# Skeletal muscle ceramides do not contribute to physical inactivity-induced insulin resistance

**DOI:** 10.1101/494559

**Authors:** Zephyra Appriou, Kévin Nay, Nicolas Pierre, Dany Saligaut, Luz Lefeuvre-Orfila, Brice Martin, Thibault Cavey, Martine Ropert, Olivier Loréal, Françoise Rannou-Bekono, Frédéric Derbré

**Affiliations:** Laboratory “Movement Sport and health Sciences”, University of Rennes -ENS Rennes, Bruz, France; GIGA-R - Translational Gastroenterology, Liège University, Belgium; INSERM NuMeCan UMR 1274, CIMIAD, France, Faculty of Medicine, University of Rennes, Rennes, France; Laboratory of Biochemistry, University Hospital Pontchaillou, Rennes, France

**Keywords:** NF-B, HOMA-IR, AMP kinase, Akt, triglycerides

## Abstract

**SUMMARY STATEMENT:** This study supports that muscle ceramide do not play a key role in insulin resistance which developed early with physical inactivity.

**ABSTRACT:** Physical inactivity increases the risk to develop type 2 diabetes, a disease characterized by a state of insulin resistance. By promoting inflammatory state, ceramides are especially recognized to alter insulin sensitivity in skeletal muscle. The present study was designed to analyze, in mice, whether muscle ceramides contribute to physical inactivity-induced insulin resistance. For this purpose, we used the wheel lock model to induce a sudden reduction of physical activity, in combination with myriocin treatment, an inhibitor of *de novo* ceramide synthesis. Mice were assigned to 3 experimental groups: voluntary wheel access group (Active), a wheel lock group (Inactive) and wheel lock group treated with myriocin (Inactive-Myr). We observed that 10 days of physical inactivity induces hyperinsulinemia and increase HOMA-IR. The muscle ceramide content were not modified by physical inactivity and myriocin. Thus, muscle ceramides do not play a role in physical inactivity-induced insulin resistance. In skeletal muscle, insulin-stimulated Akt phosphorylation and inflammatory pathway were not affected by physical inactivity whereas a reduction of GLUT4 content was observed. Based on these results, physical inactivity-induced insulin resistance seems related to a reduction in GLUT4 content rather than defects in insulin signaling. We observed in inactive mice that myriocin treatment improved glucose tolerance, insulin-stimulated Akt, AMPK activation and GLUT4 content in skeletal muscle. Such effects occur regardless of changes in muscle ceramide content. These findings highlight that myriocin could be a promising drug to improve glucose tolerance and insulin sensitivity.

## INTRODUCTION

Physical inactivity is now recognized as a global pandemic promoting the development of numerous chronic diseases including coronary heart diseases, type 2 diabetes, cancer or dementia (Booth et al., 2017; Pedersen, 2009). Each year, non-communicable chronic diseases kill 36 million people worldwide, including 17.3 million of deaths due to cardiovascular diseases and 1.3 million due to diabetes (Lee et al., 2012). Nine million of these deaths occur before 60 years old, while over 5.3 million deaths could be averted every year if all inactive people performed only 15 to 30 min/day of moderate physical exercise (Lee et al., 2012). In addition to morbidity and premature mortality, the economic burden of physical inactivity for national governments is estimated worldwide to 53.8 billion US dollars for health-care systems, and 13.7 billion US dollars for productivity losses (Ding et al., 2016).

For the World Health Organization, a person is considered as physically inactive if he doesn’t meet any of these 3 criteria: 30 min of moderate-intensity physical activity on at least 5 days every week, 20 min of vigorous-intensity physical activity on at least 3 days every week, or an equivalent combination achieving 600 metabolic equivalent (MET)-min per week (Hallal et al., 2012). Two main experimental approaches are used to understand the metabolic disorders related to this deleterious behavior (Pierre et al., 2016): 1) physical inactivity, induced by the reduction of the daily number of steps performed (from 10000 to less than 5000) in humans (Knudsen et al., 2012; Krogh-Madsen et al., 2010; Reynolds et al., 2015) and the wheel locked model in rodents (Roberts et al., 2012); 2) immobilization, induced by hindlimb unloading in rodents and bed rest in humans. Even if the latter represents an interesting approach to explore metabolic changes occurring with dramatic reduction of skeletal muscle activity (Bergouignan et al., 2011), it can be considered as too extreme compared to what physically inactive people really experience (Pierre et al., 2016; Roberts et al., 2012). In the present study, we thus choose the wheel lock model to study physical inactivity in mice.

The physiological mechanisms responsible for the development of chronic diseases related to physical inactivity has been deeply explored during the last decade (Booth et al., 2017; Gratas-Delamarche et al., 2014; Pedersen, 2009; Pierre et al., 2016). Among the proposed mechanisms, insulin resistance associated to chronic hyperinsulinemia is considered as a key triggering event promoting lipid storage and obesity (Softic et al., 2012; Stumvoll et al., 2000), but also tumor growth and cancer (Tsujimoto et al., 2017). Insulin resistance is clinically defined as the inability of a known quantity of exogenous or endogenous insulin to increase glucose uptake and utilization. In humans, several studies reported that a sudden reduction of physical activity causes whole-body insulin resistance after only few days (Knudsen et al., 2012; Krogh-Madsen et al., 2010; Reynolds et al., 2015). In this context, the drastic reduction of energy expenditure rapidly occurring in skeletal muscle is currently considered as the primary event promoting insulin resistance (Booth et al., 2017). The related gain of fat mass and the chronic low grade inflammatory state observed in inactive people are generally proposed as responsible for insulin resistance development (Booth et al., 2017; Pedersen, 2009). However, insulin resistance occurs after few days of physical inactivity whereas systemic inflammatory markers (e.g. TNF-α, IL-6) or visceral fat mass generally increase after several weeks in both humans and rodent experiments (Hamburg et al., 2007; Krogh-Madsen et al., 2010; Olsen et al., 2008; Rector et al., 2008). Thus, adipose tissue and inflammation processes do not appear to be the culprits of early insulin resistance.

Physical inactivity, whatever the experimental model, induces a shift in fuel metabolism in favor of carbohydrate oxidation and in detriment of lipid oxidation, resulting in an accumulation of intramuscular lipids (Bergouignan et al., 2006; Laye et al., 2009; Momken et al., 2011). Lipid accumulation in skeletal muscle is well known to be related with insulin resistance, especially due to their conversion into ceramides (Samuel and Shulman, 2012). Ceramides are bioactive mediators involved in cell responses to stress, and their increase in skeletal muscle is known to induce insulin resistance through the inhibitory phosphorylation of proteins of the insulin pathway including insulin receptor substrate-1/2 (IRS1/2) and phosphatidylinositol-3-kinase, (PI3-kinase) (Chavez and Summers, 2003). Ceramides are synthesized through both stimulation of sphingomyelinase-mediated hydrolysis of membrane sphingomyelin, and *de novo* synthesis pathway consisting of the condensation of palmitoyl-CoA with serine (Hannun and Luberto, 2000). Whereas accumulation of intramuscular ceramides has been extensively explored in the context of obesity (Schmitz-Peiffer, 2010; Ussher et al., 2010), their role in the onset of insulin resistance related to physical inactivity remains poorly understood.

We hypothesized that physical inactivity rapidly increases ceramide synthesis in skeletal muscle, which in turn would induce insulin resistance. Therefore, the present study was designed to analyze, in mice, whether inhibition of ceramide synthesis prevents insulin resistance observed after a short period of physical inactivity. For this purpose, we used: 1) the wheel lock model to study the effect of physical inactivity; 2) myriocin, an inhibitor of *de novo* synthesis of ceramides (Salaun et al., 2016).

## MATERIAL AND METHODS

All procedures described below were approved by the French Ministry of Higher Education and Research in accordance with the local committee on Ethics in Research of Rennes (veterinary service of health and animal protection, authorization 01259.03).

### Animal care and protocol

To mimic the effects of physical inactivity in mice, we used the wheel lock model first developed by Frank Booth’s research group (Roberts et al., 2012). Twenty-two male C57BL/6 mice were obtained at weaning (3 weeks) and allowed to acclimatize for 1 week. The animals were housed in temperature controlled room (21 ± 2°C) with a 12-h:12-h light/dark cycle and received standard rodent chow and water *ad libitum*. After 1 week, mice were separated into individual cages all equipped with a voluntary running wheel outfitted with a lap counter (IntelliBio, Nancy, France). Food intake and covered distance were daily noticed whereas body weight was recorded every week. The distance daily covered increased the first two weeks and remained unchanged during the next 4 weeks. At the end of this period, mice were assigned to 3 experimental groups with equal mean and standard deviation for daily covered distance: voluntary wheel access group (Active, n=7), a wheel lock group (Inactive, n=7) and wheel lock group treated with myriocin (Inactive-Myr, n=8). Myriocin was daily injected (0.3 mg/kg, i.p) as previously described (Hojjati et al., 2005; Lee et al., 2010). In the same schedule, Active and Inactive groups received saline vehicle. After 10 days of physical inactivity, mice were sacrificed in an overnight fasting state. Mice were anesthetized with a ketamine-xylazine-butorphanol cocktail. Adipose tissue, *rectus femoris* (RF) and right *tibialis anterior* (TA) muscles were removed, and then immediately frozen in liquid nitrogen. Left *tibialis anterior* muscles were removed, and then submitted to an *ex vivo* insulin sensitivity test. Immediately following tissue harvest, mice were euthanized via exsanguination of the heart. Intracardiac blood was collected into dry tubes and centrifuged (1,500 g, 10 min) for serum sampling.

### Glucose tolerance assessment

Oral glucose tolerance test (OGTT) was performed on the morning one day before sacrifice. Mice were fasted for 6h before the test. Then, glucose was administrated by oral gavage (1 g/kg body weight). Glucose values were obtained at rest and 15, 30, 45, 60, 90, 120 min after glucose gavage from tail blood samples. Glucose level was determined using a glucose meter (Freestyle Papillon Vision). Wheels were locked during the whole time of fasting and OGTT.

### Ex vivo muscle insulin sensitivity test

As previously described (Tardif et al., 2011), left *tibialis anterior* muscles were longitudinally divided in two strips (20-25 mg) and each strip was preincubated for 30 min in 3 mL of modified Krebs Ringer buffer (120 mM NaCl, 4.8 mM KCl, 25 mM NaHCO3, 2.5 mM CaCl2, 1.24 mM NaH2PO4, 1.25 mM MgSO4, 8 mM D-Glucose, 2 mM sodium pyruvate, 2 mM HEPES, pH 7.4) saturated with a 95% O2 and 5% CO2 mix at 37°C under stirring. One of the two strips was stimulated with 20 nM insulin for 30 minutes, then muscle samples were frozen in liquid nitrogen until analysis. Muscle insulin sensitivity was assessed by measuring the phosphorylation state of Akt on serine 473, an intermediate of the insulin pathway.

### Serum parameters

Glucose concentration was performed using Automated Beckman Coulter (Beckman Coulter, Brea, CA). Serum insulin concentrations were measured by enzyme-linked immunosorbent assay (ELISA) according to manufacturer’s instructions (Millipore, St Louis, MO, USA). Insulin sensitivity was determined by calculating HOMA-IR according to the following formula (Matthews et al., 1985): HOMA-IR = [fasting glucose (mmol/l)] × [fasting insulin (μU/ml)] /22.5.

### Quantification of muscle triglycerides

Muscle triglycerides were determined by using DiaSys kit (Diagnostic System, Grabels, France) following a preliminary organic phase extraction according to Bligh & Dyer’s method (Bligh and Dyer, 1959). Briefly, 30 mg of right *tibialis anterior* samples were crushed with 300 μL of 150 mM sodium chloride. Then 150 μL of muscle homogenates were extracted with 600 μL of a methanol-chloroform mixture (1:1, v/v). The organic layers were collected after centrifugation (10,000g for 10 min) and dried under nitrogen. Dry samples were reconstituted in 37.5 μL of isopropanol/acetonitrile/water mixture (2:1:1, v/v/v) and 10 μL were analyzed according to the manufacturer recommendations.

### Quantification of muscle sphingolipids and ceramides

Sphingolipids were extracted from ≈ 30 mg of homogenized *tibialis anterior* muscles using acidified cyclohexane/isopropanol mixture (60:40, v/v, 0.1% formic acid) and purified on NH_2_ SPE cartridges (silica gel cartridges, 100 mg) to obtain distinct fractions of ceramides and sphingomyelins (Bodennec et al., 2000). The sphingolipid fractions were then quantified by UHPLC-ESI-MS/MS using an Acquity H-Class UHPLC system (Waters, Milford, MA) combined with a Waters Xevo TQD triple quadrupole mass spectrometer. Lipid extracts were injected onto a C18 BEH column (2.1 mm x 50.0 mm, 1.7-μm particles; Waters) held to 43°C to separate all species of ceramides and sphingomyelins with two different LC elution gradients. For ceramide species separation, the gradient started at 95% of eluent B (mobile phase of water and of methanol with 1% formic acid and 5 mM ammonium formate), up to 98% in 4 min, then rapidly decreased to 95% for 0.1 min, and was maintained for 2 min. For sphingomyelin species separation, the gradient started at 95% of eluent B, up to 99% for 6 min, then rapidly decreased to 95% for 0.1 min, and was maintained for 1.4 min. Multiple reaction-monitoring mode in positive electrospray ionization was used to quantify each species of ceramides and sphingomyelins. The source heater temperature hold at 150°C, and the capillary voltage was set at 3.2 kV. The flow rate of desolvation gas was of 650 l/h at 350°C, and the cone voltage varied from 26–58 V. Argon was used as the collision gas, and collision energies varied from 12–40 eV. Data analyses were performed by Mass Lynx software version 4.1 (Waters, Manchester, UK). Sphingolipid quantification was possible with calibration curves constructed by plotting the peak area ratios of analyses to the respective internal standard against concentration using a linear regression model. The quantification measurements were performed using the TargetLinks software (Waters).

### RNA extraction and quantitative real-time PCR

Total RNA extraction from frozen *rectus femoris* was performed with Trizol^®^ (Invitrogen, France) according to the manufacturer’s instructions. The RNA quality and quantity were assessed by FlashGel DNA System^®^ and Nanodrop^®^ spectrophotometry, respectively. Reverse transcription was performed on a T100 Thermal Cycler (Bio-Rad) with iScript™ cDNA synthesis kit (Bio-Rad) from 1 μg total RNA. Real time PCR experiments were done on a CFX96 Real Time System (Bio Rad). Samples were analyzed in duplicate in 10 μl reaction volume containing 4.8 μl IQ™Sybr^®^GreenSuperMix (Bio-Rad), 0.1 μl of each primer (100 nM final) and 5 μl of diluted cDNA. The following primer sequences were used: GLUT4: (F: GCCT GCCCGAAAGAGT CT A A and R: C ATT GAT GCCT GAGAGCT GTT G); PPIA (F: CGTCTCCTTCGAGCTGTTTG and R: CC ACCCT GGC AC AT GAAT C); HPRT (F: AGGCC AGACTTT GTT GGATTT and R: CAGGACTCCTCGTATTTGCAG); RPL19 (F: CAATGCCAACTCTCGTCAACAG and R: CATCCAGGTCACCTTCTCGG). GLUT4 was normalized using three reference genes (PPPIA, HPRT, RPL19) according to geNorm analysis (Vandesompele et al., 2002).

### Western Blotting

Cytosolic protein extraction was performed from *rectus femoris* muscle in cold lysis buffer containing 10 mM Tris-HCl, pH 7.4, 0.5 M sucrose, 50 mM NaCl, 5 mM EDTA, 30 mM Na_4_P_2_O_7_, 1% NP-40, 0.25% sodium deoxycholate, 50 mM NaF, 100 μM sodium orthovanadate and proteases inhibitors cocktail (Sigma P8340, 5 μl/ml). The samples were homogenized using a Polytron homogenizer at 4°C. Each sample was then incubated on ice for 30 min followed by 3 x 10 s of sonication. The homogenates were then centrifugated at 12,000g for 12 min at 4°C. The protein concentration of the supernatant was determined by a Lowry assay using bovine serum albumin (BSA) as standard. Samples were then diluted in SDS-PAGE sample buffer [50 mM Tris-HCl, pH 6.8, 2% SDS, 10% glycerol, 5% β-mercaptoethanol, and 0.1% bromophenol blue], and heated 5 min at 95°C until analyses. Fifty micrograms of proteins were resolved on 12.5% SDS-PAGE. The proteins were transferred at 240 mA for 90 min onto a 0.2-μm nitrocellulose membrane. Membranes were blocked with 5% BSA or nonfat dry milk in TBST (Tris-buffered saline - 0.05% Tween-20) for 1 h at room temperature. Membranes were incubated overnight at 4°C with appropriate primary antibodies: AKT (1:1000, Cell Signaling), p-AKT^Ser473^ (1:1000, Cell Signaling), AMPK (1:1000, Cell Signaling), p-AMPK^Thr172^ (1:1000, Cell Signaling), p65 (1:1000, Cell Signaling), p-p65^Ser536^ (1:1000, Cell Signaling), IRS-1 (1:1000, Cell Signaling), pIRS-1^Ser302^ (1:1000, Cell Signaling), STAT3 (1:1000, Cell Signaling), p-STAT3^Ser727^, IκBα (1:1000, Cell Signaling), GLUT4 (1:1000, Abcam), and α-actin (1:700, Sigma Aldrich). Thereafter, membranes were washed with TBST and incubated for 1 h at room temperature with infrared dye-conjugated secondary antibodies (LI-COR, Lincoln, NE, USA). After washing, blots were captured using the Odyssey Imaging System (LI-COR). All blots were scanned, densitometric analysis of the bands was conducted using GS-800 Imaging densitometer and QuantityOne software. Phosphospecific signal was normalized to the total signal to estimate the ratio of activated markers.

### Statistical analysis

Data are presented as mean ± SEM. Normality and equality of variances were checked using a Kolmogorov-Smirnov and Fischer test, respectively. A one-way analysis of variance (ANOVA) was performed to compare each parameter between the 3 experimental groups. When appropriate, the Fisher LSD test was used as a post-hoc analysis. If normality and/or equal variance tests failed, we checked the significance using one-way ANOVA on ranks (Kruskal-Wallis). When appropriate, the Dunn’s test was used as a post-hoc analysis. For all statistical analyses, the significance level was set at 0.05. Data were analyzed using the statistical package GraphPad Prism version 6.02 for Windows (GraphPad Software, La Jolla, California).

## RESULTS

### Physical activity levels, body weight, food intake and visceral fat mass

During the 10 days of wheel lock, active mice exhibited a mean daily physical activity levels of 3.90 ± 0.78 km/day. Daily food intake was lower in both Inactive and Inactive-Myr mice compared to Active mice (−7.9% and −7.6%, p=0.007 and 0.01, respectively, Table 1). After 10 days of wheel lock, body weight significantly increased only in Active mice. No significant difference of body weight was observed between the 3 experimental groups before and after the 10 days of physical inactivity. Visceral fat mass did not differ between the 3 experimental groups at the end of the protocol (Table 1).

**Table 1.** Effects of physical inactivity and myriocin treatment on body weight, food intake and visceral fat mass. Data are presented as mean ± SEM. Significant differences (p< 0.05) are indicated as follows: comparison vs. values before wheel locking ($), comparison vs. Active group (*)

|  | Active | Inactive | Inactive-Myr |
| --- | --- | --- | --- |
| **Body weight (g)** |  |  |  |
| **Before locking** | 25.2 ± 0.5 | 25.8 ± 0.2 | 24.8 ± 0.2 |
| **10 days post-locking** | 25.8 ± 0.6<sup>\$</sup> | 25.9 ± 0.2 | 24.5 ± 0.3 |
| **Food intake (g/day)** | 4.16 ± 0.05 | 3.83 ± 0.09<sup>*</sup> | 3.84 ± 0.08<sup>*</sup> |
| **Physical activity (km/day)** | 3.90 ± 0.78 | ----- | ----- |
| **Visceral fat mass (g/100 g BW)** | 2.12 ± 0.19 | 2.30 ± 0.25 | 2.08 ± 0.15 |

### Muscle triglycerides and ceramides content

We first observed that muscle triglyceride (TG) content tend to increase in Inactive mice (p=0.09, Fig. 1A) whereas it increase in the Inactive-Myr group (p=0.027, Fig. 1A). Contrary to our hypothesis, physical inactivity did not modify total, saturated and unsaturated ceramides content in muscle (Fig. 1B). Individual ceramides species ranging from C16:0 to C24:1 remained also unchanged after 10 days of physical inactivity (Fig. 1C and 1D). Surprisingly, muscle ceramides content were not affected by 10 days of myriocin treatment (Fig. 1B, 1C and 1D).

**Figure 1.**
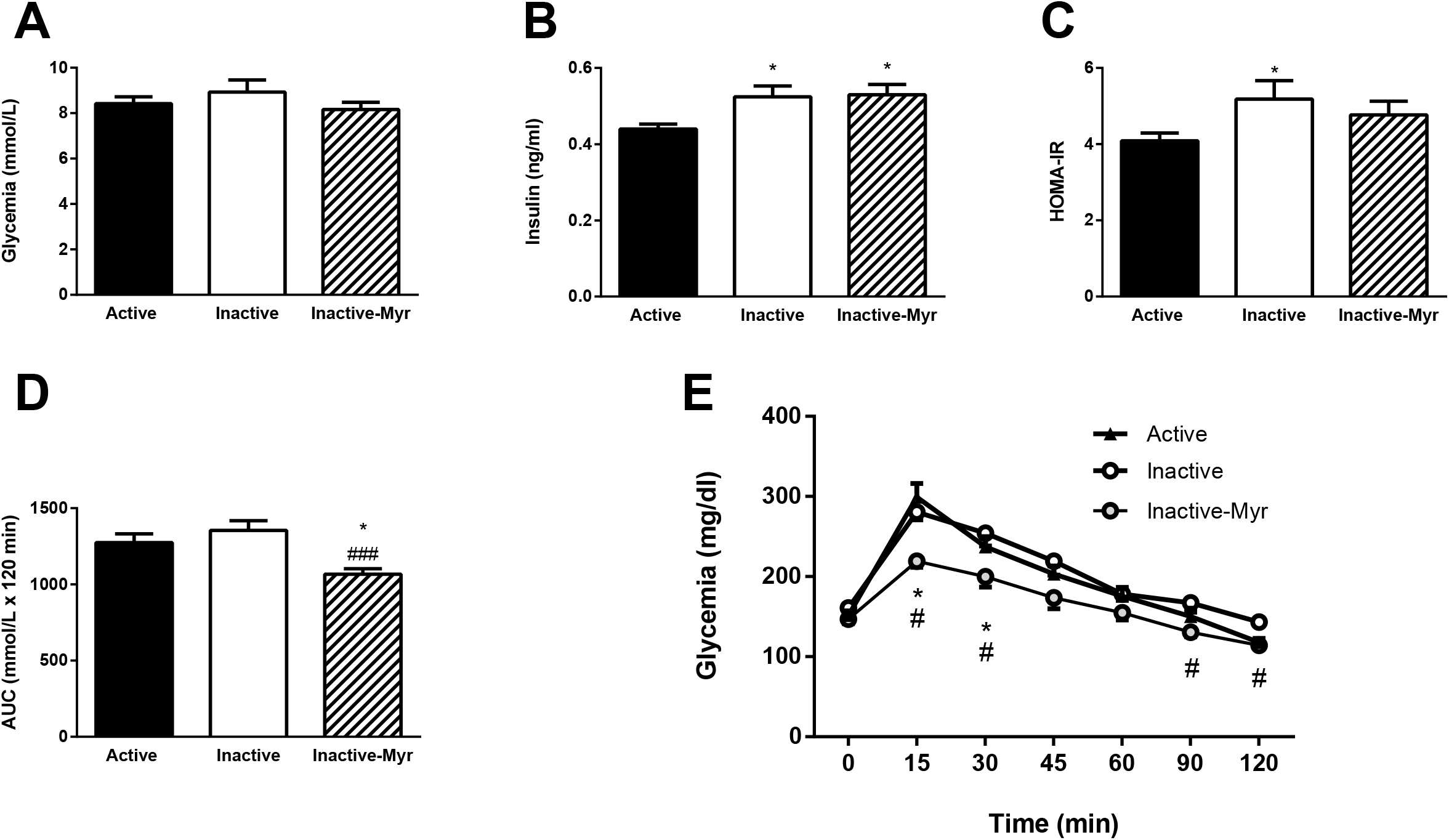
Effects of physical inactivity and myriocin treatment on lipids profile. Data are presented as mean ± SEM (n=7-8/group). *A*: triglycerides content. *B*: total, saturated, and unsaturated ceramide content. *C*: unsaturated species profile. *D*: saturated species profile. Significant differences are indicated as follows: comparison vs. Active group (*: p<0.05)

### Whole-body insulin sensitivity and glucose tolerance

The effect physical inactivity and myriocin treatment on whole-body insulin sensitivity was assessed by measuring the circulating level of glucose and insulin. Fasting glycemia remained unaffected with physical inactivity and myriocin treatment (Fig. 2A). Serum insulin levels were higher in both Inactive and Inactive-Myr groups compared to Active group (p=0.025 and 0.015, respectively, Fig. 2B). Such results were associated with a higher HOMA-IR index in Inactive compared to Active mice (p=0.045, Fig. 2C), whereas this index did not differ between Inactive-Myr and Active mice (p=0.17, Fig. 2C). Thus, myriocin seems to prevent physical inactivity-induced whole-body insulin resistance. An OGTT was also performed at the end of the protocol to observe the effects of physical inactivity and myriocin on glucose tolerance. Here, we did not observe significant difference in glucose concentrations (at all time points) and area under the curve (AUC) between Active and Inactive mice (Fig. 2D and E). However, we reported that Inactive-Myr mice exhibited an AUC significantly lower compared to both Active (p=0.019, Fig. 2D) and Inactive mice (p<0.001, Fig. 2D). This effect was due to a lower glucose levels both at the beginning and the end of OGTT (15 min, 30 min, 90 and 120 min, p<0.05, Fig. 2E). Taken together, our results indicate that myriocin improves glucose tolerance.

**Figure 2.**
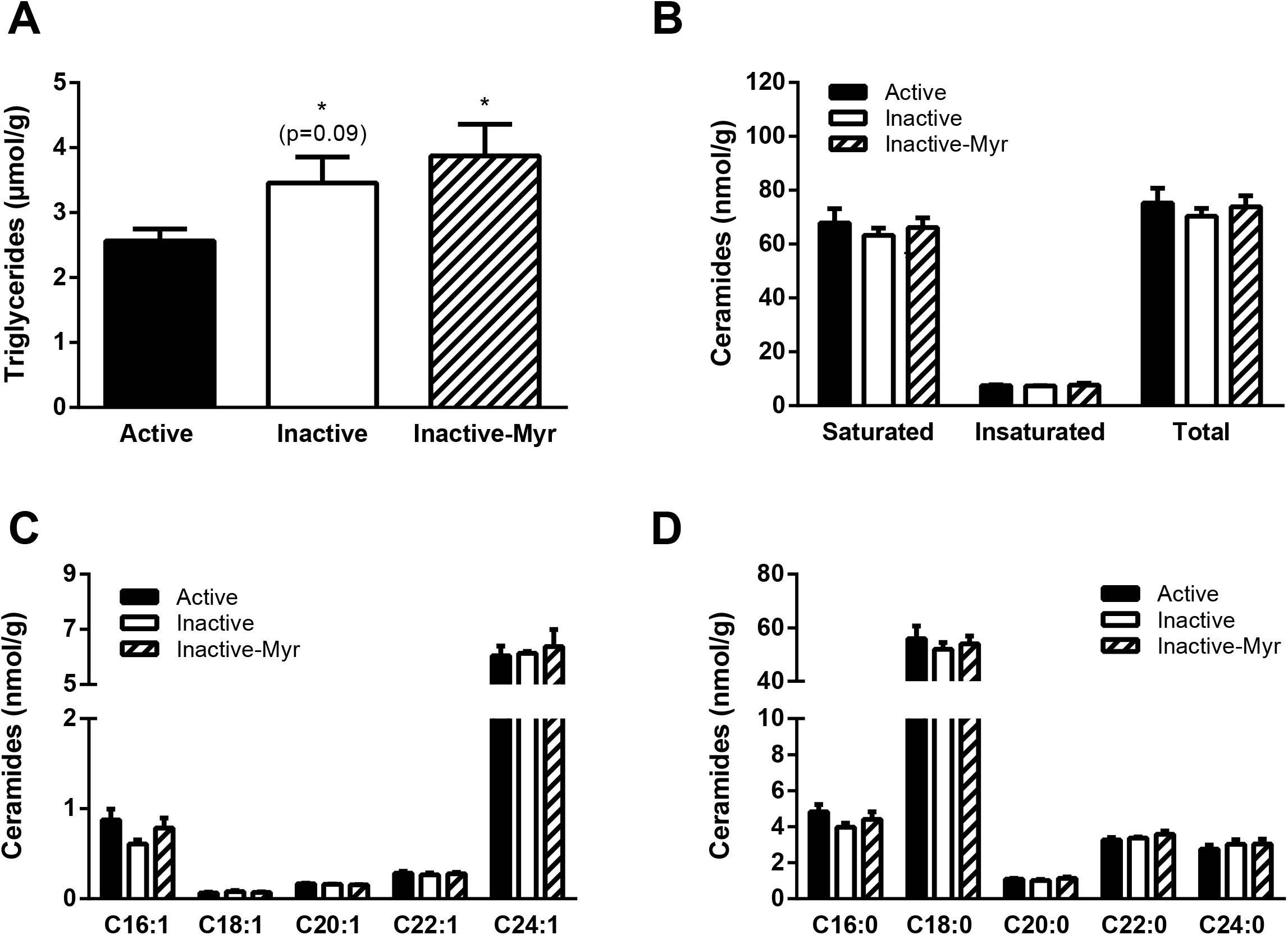
Effects of physical inactivity and myriocin treatment on whole-body insulin sensitivity and glucose tolerance. Data are presented as mean ± SEM (n=7-8/group). *A*: serum glucose concentrations. *B*: serum insulin concentrations. *C*: HOMA-IR index. *D*: area under the curve (AUC) for OGTT. *E*: evolution of glycemia following oral glucose load. Significant differences are indicated as follows: comparison vs. Active group (*: p<0.05), comparison vs. Inactive group (###: p<0.001)

### Proximal insulin signaling and GLUT4 content in skeletal muscle

We observed a deleterious effect of physical inactivity on whole-body insulin sensitivity and a beneficial effect of myriocin on glucose tolerance. To investigate the mechanism regulating such effects, we first evaluated the integrity of insulin signaling in skeletal muscle by measuring, *ex vivo*, basal and insulin-stimulated Akt activation. We observed that insulin stimulation increased phospho-Akt levels in Active (p=0.019), Inactive (p=0.029) and Inactive-Myr (p<0.001, Fig. 3A). Whereas the magnitude of these responses did not differ between Active and Inactive mice, Inactive-Myr mice exhibited a higher muscle activation of Akt in response to insulin when compared to both Active and Inactive mice (p=0.046 and p=0.036, respectively, Fig. 3A). Glucose uptake is recognized to be regulated by insulin signaling and GLUT4 pool in skeletal muscle (Pierre et al., 2016). In the present study, we observed that physical inactivity reduced GLUT4 protein (p=0.043, Fig. 3B and 3E), but not GLUT4 mRNA (Fig. 3C). Interestingly, we observed that myriocin treatment prevent the reduction of muscle GLUT4 protein induced by physical inactivity (p=0.005, Fig. 3B). At the mRNA level, GLUT4 was reduced by myriocin in Inactive-Myr compared to Inactive mice (p=0.031, Fig. 3B). As AMP-activated protein kinase (AMPK) is recognized to regulate GLUT4 expression and it translocation to the membrane (McGee et al., 2008), we decided to measure the levels of AMPK activation in skeletal muscle. We reported that AMPK activation remained unchanged in Inactive compared to Active mice (Fig. 3D and E). Interestingly, we showed that AMPK activation was higher in myriocin-treated mice compared to both Active and Inactive mice (p=0.031 and p=0.003, respectively, Fig. 3D and 3E). In insulin-resistant rodent models, chronic hyperphosphorylation of IRS1 on Ser^302^ has been identified as playing a key role in muscle insulin resistance (Morino et al., 2008). In our experiments, no significant change was observed in the phosphorylation state of IRS1 on Ser^302^ between the 3 experimental groups (Fig. 4A and 4E). Muscle insulin resistance is also related to activation of NF-KB and IL6/STAT3 inflammatory signaling pathways (Gratas-Delamarche et al., 2014). Here, we reported that physical inactivity combined or not with myriocin treatment did not modulate phospho-p65 and IKBα (Fig. 4C and 4E), two well-recognized markers of NF-KB activation (Christian et al., 2016). Similarly, phosphorylation of STAT3 on Ser^727^ remained unchanged in the three experimental groups (Fig. 4B and 4E).

**Figure 3.**
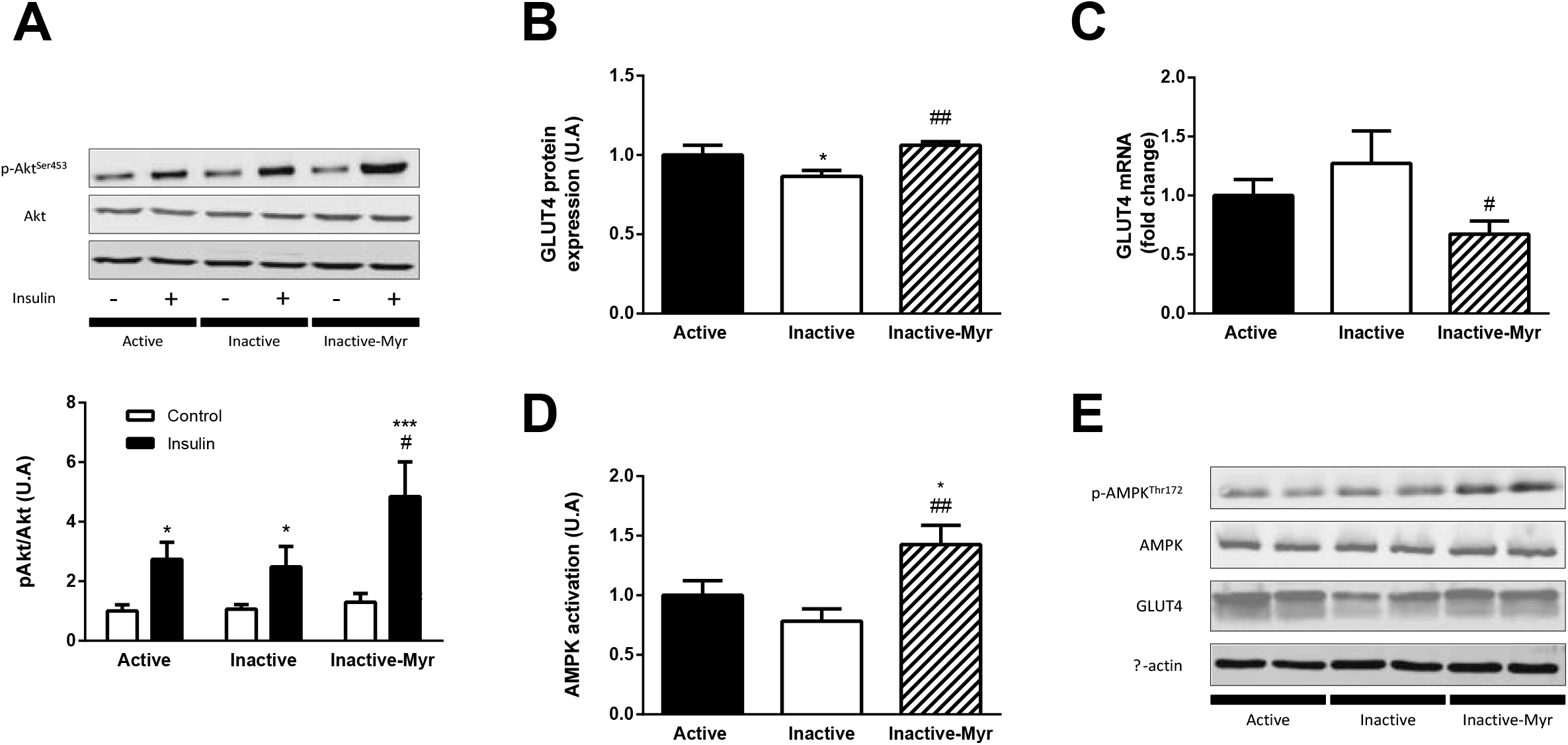
Effects of physical inactivity and myriocin treatment on muscle insulin signaling. Data are presented as mean ± SEM (n=7-8/group). *A*: Insulin-stimulated Akt activation in skeletal muscle of different experimental groups. *B*: muscle GLUT4 protein content muscle. *C*: GLUT4 mRNA levels. *D*: muscle AMPK activation. *E*: representative Western Blot of the data obtained in panel C and D. Significant differences are indicated as follows: comparison vs. Active group (*: p<0.05), comparison vs. Inactive group (#: p<0.05, ##: p<0.01)

**Figure 4.**
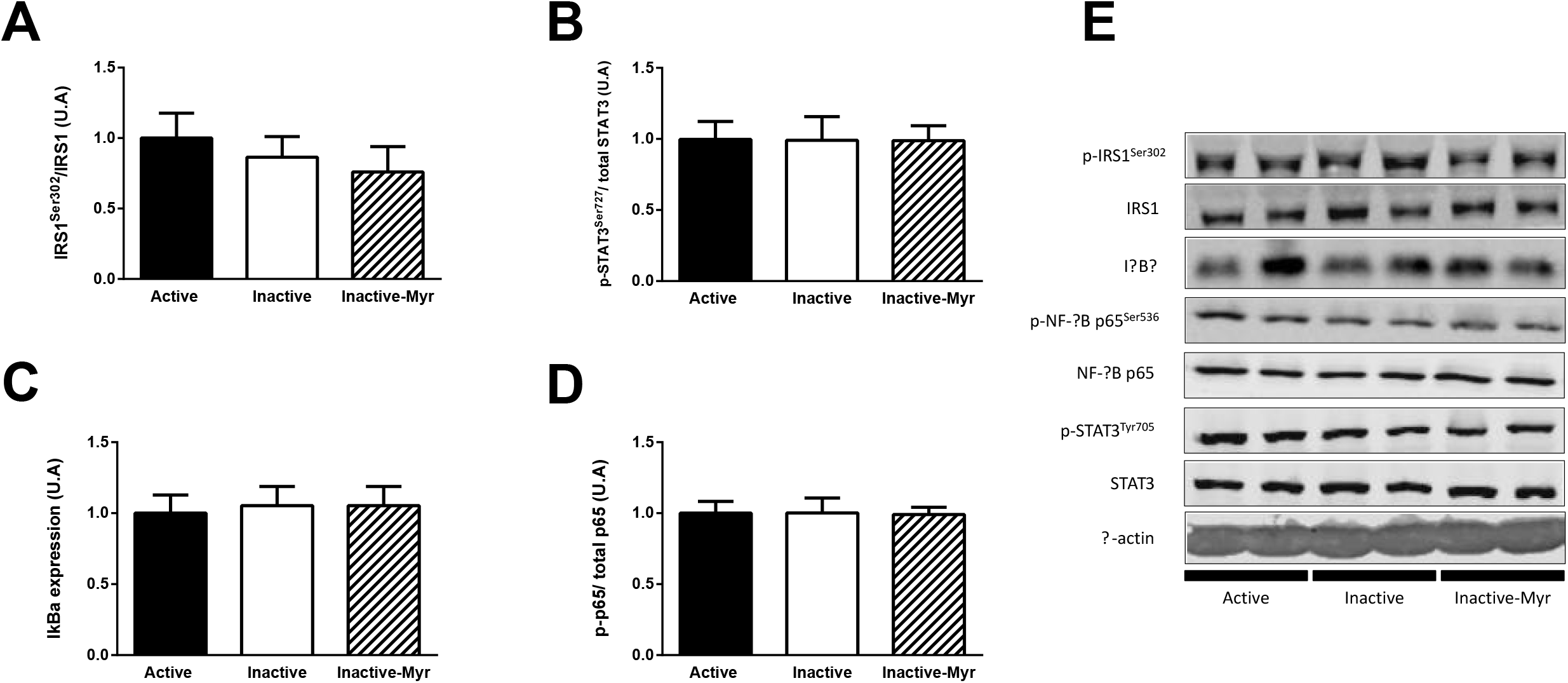
Effects of physical inactivity and myriocin treatment on signaling pathways involved in muscle insulin resistance. Data are presented as mean ± SEM (n=7-8/group). *A*: IRS1^Ser302^ activation. *B*: STAT3 activation. *C*: IκBα content. *D*: NF-κB p65 activation. *E*: representative Western Blot analyses of proteins involved in muscle insulin resistance.

## DISCUSSION

Muscle ceramides are known to play a key role in the development of insulin resistance during high fat nutritional intake, but their role in physical inactivity-induced insulin resistance are poorly understood. Our data support that early insulin resistance observed with physical inactivity is not due to ceramide accumulation but could be caused by a reduction in muscle GLUT4 content. We also observed that 10 days of myriocin treatment in inactive mice improve glucose tolerance, but surprisingly through a mechanism that appears independent of changes in muscle ceramide content.

Immobilization rapidly causes both in humans and rodents a reduction in fatty acid (FA) transport to mitochondria and in mitochondrial FA β-oxidation, resulting in an accumulation of intramuscular lipids (Bergouignan et al., 2006; Kwon et al., 2016; Laye et al., 2009; Momken et al., 2011; Salaun et al., 2016). Saturated FA oxidation, including palmitate, are particularly reduced in skeletal muscle during immobilization (Bergouignan et al., 2011). Interestingly, palmitate is well identified to increase the expression of serine palmitoyltransferase 2 (SPT2), a key enzyme in ceramide biosynthesis (Erickson et al., 2012). Ceramides have been identified as key bioactive sphingolipids in the development of muscle insulin resistance, especially the C:16 and C:18 moieties, which are the most abundant in skeletal muscle (Chung et al., 2017; Perreault et al., 2018). Based on previous data from our laboratory and others showing in rodents that C:16 and C:18 ceramides increase in skeletal muscle after 1 or 2 weeks of immobilization (Kwon et al., 2016; Salaun et al., 2016), we hypothesized that 10 days of physical inactivity induced by wheel lock will also cause muscle ceramide accumulation. Herein, we show that muscle ceramides are not responsible of insulin resistance induced by physical inactivity. Contrary to immobilization, we observed that total and saturated ceramide levels in muscle remained unchanged after 10 days of physical inactivity. This was also the case for muscle TG which are increase after immobilization (3, 50), while we reported only a trend with physical inactivity. All together, these results support that immobilization and physical inactivity are not equivalent models to study the effect of a reduction of energy expenditure.

In rodents, the wheel lock model is currently the closest model to mimic human physical inactivity (Roberts et al., 2012). Using this model in mice, we observed that 10 days of physical inactivity are sufficient to induce hyperinsulinemia and to increase the HOMA-IR, both considered as hallmarks of insulin resistance (Singh and Saxena, 2010). These results are in accordance with previous studies conducted in Sprague-Dawley or OLETF rats after 7 days of wheel lock (Rector et al., 2010, 2008; Teich et al., 2017). Few data are available about the effects of the wheel lock model on glucose tolerance. Teich and colleagues recently observed that 7 days of wheel lock were sufficient to affect glucose tolerance in young Sprague-Dawley male rats (Teich et al., 2017). In the present study, we did not report an effect on glucose tolerance after 10 days of wheel lock. Our results are in accordance with data obtained in humans exposed to a reduction in daily number of steps during 5 or 14 days, where no change in glucose tolerance was reported (Knudsen et al., 2012; Reynolds et al., 2015). All together, these results sustain that the wheel lock model is a realistic experimental set-up to study, in mice, human physical inactivity and its role in chronic diseases development.

Skeletal muscle being responsible of 80% of whole-body glucose uptake under insulin stimulation (DeFronzo et al., 1985), early insulin resistance occurring with physical inactivity is mainly attributed to alterations in skeletal muscle glucose uptake (Knudsen et al., 2012; Krogh-Madsen et al., 2010). Interestingly, using the wheel lock model during only 2 days, Kump and Booth observed a reduction of insulin-stimulated 2-deoxyglucose uptake in skeletal muscle (Kump and Booth, 2005). Similar results were also reported in rats submitted to muscle unloading after 24h (Kawamoto et al., 2016; O’keefe et al., 2004). However, the mechanisms by which physical inactivity and muscle unloading rapidly induce muscle insulin resistance remain unclear. Krogh-Madsen and colleagues (2010) have reported that a reduction of muscle Akt phosphorylation during hyperinsulinemic euglycemic clamp occurred after 2 weeks of reduction of ambulatory activity, thus supporting the idea that insulin signaling was early affected by physical inactivity. However, if the clamp is the gold standard to assess peripheral insulin sensitivity, it appears less appropriate to assess insulin signaling. Indeed, presence of circulating hormones or cytokines (e.g. leptin, adiponectin, IL-6) and muscle contractility are all factors affecting muscle insulin signaling during clamp, independently of direct action of insulin. For these reasons, we decided to explore insulin-stimulated muscle Akt phosphorylation in *ex-vivo* conditions to determine whether insulin signaling was affected with a short period of physical inactivity. In this condition, we observed that Akt phosphorylation in inactive mice did not differ from active animals, suggesting that insulin signaling until Akt remains unaffected after 10 days of physical inactivity. To support such results, we explored the effects of physical inactivity on inflammatory pathways recognized to affect insulin signaling. Here, neither NF-κB nor IL-6/STAT3 signaling pathways were affected in skeletal muscle of inactive mice. Further, in response to pro-inflammatory state IRS1 is inhibited by a phosphorylation on its Ser^302^ (Gratas-Delamarche et al., 2014; Hage Hassan et al., 2016; Morino et al., 2008). In the present study, we found that physical inactivity did not modify the phosphorylation state IRS1^Ser302^. Taken together, these results indicate that, in the context of physical inactivity, the onset of insulin resistance is not due an inflammatory process affecting the skeletal muscle.

Our data support that muscle insulin signaling does not appear affected by a short period of physical inactivity. In accordance with Kump and Booth (2005), we reported that 10 days of wheel lock caused a significant reduction of GLUT4 pool in skeletal muscle. These results support the idea that inactivity-induced muscle insulin resistance would be rather related to a decrease of muscle GLUT4 pool than a defect in insulin signaling. Interestingly, we observed that mRNA coding for GLUT4 was not affected suggesting that the decrease of GLUT4 pool would be due to an increase of its degradation. It is well established that chronic insulin stimulation causes a decrease of GLUT4 content in adipocytes, mainly due to an accelerated GLUT4 degradation in the lysosomes (Liu et al., 2007; Ma et al., 2014, p. 4; Sargeant and Pâquet, 1993). Similar cellular events could occur in skeletal muscle of inactive. Such mechanisms need to be explored in further experiments.

To explore the role of ceramides in physical inactivity-induced insulin resistance, we treated inactive mice with myriocin, an inhibitor of *de novo* synthesis of ceramides. As previously reported in others experimental models including Zucker diabetic rats or high fat diet-fed mice (Holland et al., 2007; Ussher et al., 2010), we observed that myriocin treatment improved glucose tolerance in inactive mice. Interestingly, we also reported that myriocin treatment caused a significant increase of AMPK phosphorylation. This result is in accordance with the data of Liu and colleagues (2013) demonstrating that myriocin prolonged yeast lifespan by activating key signaling pathways controlling stress resistance and energy metabolic homeostasis including AMPK signaling pathway. Thus, the beneficial effect of myriocin on glucose tolerance could be mediated by AMPK (Jensen et al., 2014). Interestingly, we also observed higher insulin-stimulated Akt phosphorylation in myriocin-treated compared to non-treated inactive mice. Such improvement of muscle insulin signaling could be linked to AMPK activation since this kinase is recognized to enhance the response of Akt phosphorylation on serine 473 residue through the modulation of MTORC2 complex activity (Kleinert et al., 2016; Sarbassov et al., 2005). We also observed that myriocin prevents the decrease of GLUT4 induced by physical inactivity. Although AMPK has been proposed to stimulate GLUT4 transcription (Gong et al., 2011; McGee et al., 2008), we observed a decrease of GLUT4 mRNA associated with AMPK activation in Inactive-My compared to Inactive mice. Consequently, an AMPK-independent mechanism could be responsible of the effect of myriocin on muscle GLUT4 protein. To sum up, the effect of myriocin on glucose tolerance seems implicated a coordination of AMPK, Akt and GLUT4 in a manner that needs to be elucidated. Contrary to our hypothesis, the benefits of myriocin treatment on glucose tolerance are not associated to modulation in muscle ceramide concentrations. The previous studies reporting in mice a beneficial effect of myriocin on glucose tolerance and insulin sensitivity were associated to a prevention of muscle ceramide accumulation induced by high fat diet or pathological genetic background (Holland et al., 2007; Ussher et al., 2010; Yang et al., 2009). Herein, we found that myriocin and physical inactivity act on insulin sensitivity independently from ceramides, thus highlighting the need to explore other hypothesis.

In summary, our results support that muscle ceramide accumulation and inflammatory pathways do not play a role in physical inactivity-induced insulin resistance. The insulin resistance observed after a short period of physical inactivity seems related to a reduction in muscle GLUT4 content rather than to defects of muscle insulin signaling. Our findings also support that immobilization is not what inactive people experience, and that biological effects obtained with this kind of models should not associated with physical inactivity. Finally, our data highlight that myriocin could be a promising molecule to improve glucose tolerance and muscle insulin sensitivity.

## ACKNOWLEDGMENTS

The authors thank Véronique Ferchaud-Roucher and Mickael Croyal (Corsaire platform, CRNH, UMR1280, Nantes) for technical help in sphingolipid analyses.

## REFERENCES

Bergouignan, A., Rudwill, F., Simon, C., Blanc, S., 2011. Physical inactivity as the culprit of metabolic inflexibility: evidence from bed-rest studies. J. Appl. Physiol. Bethesda Md 1985 111, 1201–1210. https://doi.org/10.1152/japplphysiol.00698.2011

Bergouignan, A., Schoeller, D.A., Normand, S., Gauquelin-Koch, G., Laville, M., Shriver, T., Desage, M., Le Maho, Y., Ohshima, H., Gharib, C., Blanc, S., 2006. Effect of physical inactivity on the oxidation of saturated and monounsaturated dietary Fatty acids: results of a randomized trial. PLoS Clin. Trials 1, e27. https://doi.org/10.1371/journal.pctr.0010027

Bergouignan, A., Trudel, G., Simon, C., Chopard, A., Schoeller, D.A., Momken, I., Votruba, S.B., Desage, M., Burdge, G.C., Gauquelin-Koch, G., Normand, S., Blanc, S., 2009. Physical inactivity differentially alters dietary oleate and palmitate trafficking. Diabetes 58, 367–376. https://doi.org/10.2337/db08-0263

Bligh, E.G., Dyer, W.J., 1959. A rapid method of total lipid extraction and purification. Can. J. Biochem. Physiol. 37, 911–917. https://doi.org/10.1139/o59-099

Bodennec, J., Koul, O., Aguado, I., Brichon, G., Zwingelstein, G., Portoukalian, J., 2000. A procedure for fractionation of sphingolipid classes by solid-phase extraction on aminopropyl cartridges. J. Lipid Res. 41, 1524–1531.

Booth, F.W., Roberts, C.K., Thyfault, J.P., Ruegsegger, G.N., Toedebusch, R.G., 2017. Role of Inactivity in Chronic Diseases: Evolutionary Insight and Pathophysiological Mechanisms. Physiol. Rev. 97, 1351–1402. https://doi.org/10.1152/physrev.00019.2016

Chavez, J.A., Summers, S. A., 2003. Characterizing the effects of saturated fatty acids on insulin signaling and ceramide and diacylglycerol accumulation in 3T3-L1 adipocytes and C2C12 myotubes. Arch. Biochem. Biophys. 419, 101–109.

Christian, F., Smith, E.L., Carmody, R.J., 2016. The Regulation of NF-κB Subunits by Phosphorylation. Cells 5. https://doi.org/10.3390/cells5010012

Chung, J.O., Koutsari, C., Blachnio-Zabielska, A.U., Hames, K.C., Jensen, M.D., 2017. Intramyocellular Ceramides: Subcellular Concentrations and Fractional De Novo Synthesis in Postabsorptive Humans. Diabetes 66, 2082–2091. https://doi.org/10.2337/db17-0082

DeFronzo, R.A., Gunnarsson, R., Björkman, O., Olsson, M., Wahren, J., 1985. Effects of insulin on peripheral and splanchnic glucose metabolism in noninsulin-dependent (type II) diabetes mellitus. J. Clin. Invest. 76, 149–155. https://doi.org/10.1172/JCI111938

Ding, D., Lawson, K.D., Kolbe-Alexander, T.L., Finkelstein, E.A., Katzmarzyk, P.T., van Mechelen, W., Pratt, M., Lancet Physical Activity Series 2 Executive Committee, 2016. The economic burden of physical inactivity: a global analysis of major non-communicable diseases. Lancet Lond. Engl. 388, 1311–1324. https://doi.org/10.1016/S0140-6736(16)30383-X

Erickson, K.A., Smith, M.E., Anthonymuthu, T.S., Evanson, M.J., Brassfield, E.S., Hodson, A.E., Bressler, M.A., Tucker, B.J., Thatcher, M.O., Prince, J.T., Hancock, C.R., Bikman, B.T., 2012. AICAR inhibits ceramide biosynthesis in skeletal muscle. Diabetol. Metab. Syndr. 4, 45. https://doi.org/10.1186/1758-5996-4-45

Gong, H., Xie, J., Zhang, N., Yao, L., Zhang, Y., 2011. MEF2A binding to the Glut4 promoter occurs via an AMPKα2-dependent mechanism. Med. Sci. Sports Exerc. 43, 1441–1450. https://doi.org/10.1249/MSS.0b013e31820f6093

Gratas-Delamarche, A., Derbré, F., Vincent, S., Cillard, J., 2014. Physical inactivity, insulin resistance, and the oxidative-inflammatory loop. Free Radic. Res. 48, 93–108. https://doi.org/10.3109/10715762.2013.847528

Hage Hassan, R., Pacheco de Sousa, A.C., Mahfouz, R., Hainault, I., Blachnio-Zabielska, A., Bourron, O., Koskas, F., Górski, J., Ferré, P., Foufelle, F., Hajduch, E., 2016. Sustained Action of Ceramide on the Insulin Signaling Pathway in Muscle Cells: IMPLICATION OF THE DOUBLE-STRANDED RNA-ACTIVATED PROTEIN KINASE. J. Biol. Chem. 291, 3019–3029. https://doi.org/10.1074/jbc.M115.686949

Hallal, P.C., Andersen, L.B., Bull, F.C., Guthold, R., Haskell, W., Ekelund, U., Lancet Physical Activity Series Working Group, 2012. Global physical activity levels: surveillance progress, pitfalls, and prospects. Lancet Lond. Engl. 380, 247–257. https://doi.org/10.1016/S0140-6736(12)60646-1

Hamburg, N.M., McMackin, C.J., Huang, A.L., Shenouda, S.M., Widlansky, M.E., Schulz, E., Gokce, N., Ruderman, N.B., Keaney, J.F., Vita, J.A., 2007. Physical inactivity rapidly induces insulin resistance and microvascular dysfunction in healthy volunteers. Arterioscler. Thromb. Vasc. Biol. 27, 2650–2656. https://doi.org/10.1161/ATVBAHA.107.153288

Hannun, Y.A., Luberto, C., 2000. Ceramide in the eukaryotic stress response. Trends Cell Biol. 10, 73–80.

Hojjati, M.R., Li, Z., Zhou, H., Tang, S., Huan, C., Ooi, E., Lu, S., Jiang, X.-C., 2005. Effect of myriocin on plasma sphingolipid metabolism and atherosclerosis in apoE-deficient mice. J. Biol. Chem. 280, 10284–10289. https://doi.org/10.1074/jbc.M412348200

Holland, W.L., Brozinick, J.T., Wang, L.-P., Hawkins, E.D., Sargent, K.M., Liu, Y., Narra, K., Hoehn, K.L., Knotts, T.A., Siesky, A., Nelson, D.H., Karathanasis, S.K., Fontenot, G.K., Birnbaum, M.J., Summers, S.A., 2007. Inhibition of Ceramide Synthesis Ameliorates Glucocorticoid-, Saturated-Fat-, and Obesity-Induced Insulin Resistance. Cell Metab. 5, 167–179. https://doi.org/10.1016/j.cmet.2007.01.002

Jensen, T.E., Sylow, L., Rose, A.J., Madsen, A.B., Angin, Y., Maarbjerg, S.J., Richter, E.A., 2014. Contraction-stimulated glucose transport in muscle is controlled by AMPK and mechanical stress but not sarcoplasmatic reticulum Ca2+ release. Mol. Metab. 3, 742–753. https://doi.org/10.1016/j.molmet.2014.07.005

Kawamoto, E., Koshinaka, K., Yoshimura, T., Masuda, H., Kawanaka, K., 2016. Immobilization rapidly induces muscle insulin resistance together with the activation of MAPKs (JNK and p38) and impairment of AS160 phosphorylation. Physiol. Rep. 4. https://doi.org/10.14814/phy2.12876

Kleinert, M., Parker, B.L., Chaudhuri, R., Fazakerley, D.J., Serup, A., Thomas, K.C., Krycer, J.R., Sylow, L., Fritzen, A.M., Hoffman, N.J., Jeppesen, J., Schjerling, P., Ruegg, M.A., Kiens, B., James, D.E., Richter, E.A., 2016. mTORC2 and AMPK differentially regulate muscle triglyceride content via Perilipin 3. Mol. Metab. 5, 646–655. https://doi.org/10.1016/j.molmet.2016.06.007

Knudsen, S.H., Hansen, L.S., Pedersen, M., Dejgaard, T., Hansen, J., Hall, G.V., Thomsen, C., Solomon, T.P.J., Pedersen, B.K., Krogh-Madsen, R., 2012. Changes in insulin sensitivity precede changes in body composition during 14 days of step reduction combined with overfeeding in healthy young men. J. Appl. Physiol. Bethesda Md 1985 113, 7–15. https://doi.org/10.1152/japplphysiol.00189.2011

Krogh-Madsen, R., Thyfault, J.P., Broholm, C., Mortensen, O.H., Olsen, R.H., Mounier, R., Plomgaard, P., van Hall, G., Booth, F.W., Pedersen, B.K., 2010. A 2-wk reduction of ambulatory activity attenuates peripheral insulin sensitivity. J. Appl. Physiol. Bethesda Md 1985 108, 1034–1040. https://doi.org/10.1152/japplphysiol.00977.2009

Kump, D.S., Booth, F.W., 2005. Alterations in insulin receptor signalling in the rat epitrochlearis muscle upon cessation of voluntary exercise. J. Physiol. 562, 829–838. https://doi.org/10.1113/jphysiol.2004.073593

Kwon, O.S., Nelson, D.S., Barrows, K.M., O’Connell, R.M., Drummond, M.J., 2016. Intramyocellular ceramides and skeletal muscle mitochondrial respiration are partially regulated by Toll-like receptor 4 during hindlimb unloading. Am. J. Physiol. Regul. Integr. Comp. Physiol. 311, R879–R887. https://doi.org/10.1152/ajpregu.00253.2016

Laye, M.J., Rector, R.S., Borengasser, S.J., Naples, S.P., Uptergrove, G.M., Ibdah, J.A., Booth, F.W., Thyfault, J.P., 2009. Cessation of daily wheel running differentially alters fat oxidation capacity in liver, muscle, and adipose tissue. J. Appl. Physiol. Bethesda Md 1985 106, 161–168. https://doi.org/10.1152/japplphysiol.91186.2008

Lee, B.J., Kim, Jae Seon, Kim, B.K., Jung, S.J., Joo, M.K., Hong, S.G., Kim, Jang Soo, Kim, J.H., Yeon, J.E., Park, J.-J., Byun, K.S., Bak, Y.-T., Yoo, H.-S., Oh, S., 2010. Effects of sphingolipid synthesis inhibition on cholesterol gallstone formation in C57BL/6J mice. J. Gastroenterol. Hepatol. 25, 1105–1110. https://doi.org/10.1111/j.1440-1746.2010.06246.x

Lee, I.-M., Shiroma, E.J., Lobelo, F., Puska, P., Blair, S.N., Katzmarzyk, P.T., Lancet Physical Activity Series Working Group, 2012. Effect of physical inactivity on major non-communicable diseases worldwide: an analysis of burden of disease and life expectancy. Lancet Lond. Engl. 380, 219–229. https://doi.org/10.1016/S0140-6736(12)61031-9

Liu, J., Huang, X., Withers, B.R., Blalock, E., Liu, K., Dickson, R.C., 2013. Reducing Sphingolipid Synthesis Orchestrates Global Changes to Extend Yeast Lifespan. Aging Cell 12, 833–841. https://doi.org/10.1111/acel.12107

Liu, L.-B., Omata, W., Kojima, I., Shibata, H., 2007. The SUMO conjugating enzyme Ubc9 is a regulator of GLUT4 turnover and targeting to the insulin-responsive storage compartment in 3T3-L1 adipocytes. Diabetes 56, 1977–1985. https://doi.org/10.2337/db06-1100

Ma, J., Nakagawa, Y., Kojima, I., Shibata, H., 2014. Prolonged insulin stimulation down-regulates GLUT4 through oxidative stress-mediated retromer inhibition by a protein kinase CK2-dependent mechanism in 3T3-L1 adipocytes. J. Biol. Chem. 289, 133–142. https://doi.org/10.1074/jbc.M113.533240

Matthews, D.R., Hosker, J.P., Rudenski, A.S., Naylor, B.A., Treacher, D.F., Turner, R.C., 1985. Homeostasis model assessment: insulin resistance and beta-cell function from fasting plasma glucose and insulin concentrations in man. Diabetologia 28, 412–419.

McGee, S.L., van Denderen, B.J.W., Howlett, K.F., Mollica, J., Schertzer, J.D., Kemp, B.E., Hargreaves, M., 2008. AMP-activated protein kinase regulates GLUT4 transcription by phosphorylating histone deacetylase 5. Diabetes 57, 860–867. https://doi.org/10.2337/db07-0843

Momken, I., Stevens, L., Bergouignan, A., Desplanches, D., Rudwill, F., Chery, I., Zahariev, A., Zahn, S., Stein, T.P., Sebedio, J.L., Pujos-Guillot, E., Falempin, M., Simon, C., Coxam, V., Andrianjafiniony, T., Gauquelin-Koch, G., Picquet, F., Blanc, S., 2011. Resveratrol prevents the wasting disorders of mechanical unloading by acting as a physical exercise mimetic in the rat. FASEB J. Off. Publ. Fed. Am. Soc. Exp. Biol. 25, 3646–3660. https://doi.org/10.1096/fj.10-177295

Morino, K., Neschen, S., Bilz, S., Sono, S., Tsirigotis, D., Reznick, R.M., Moore, I., Nagai, Y., Samuel, V., Sebastian, D., White, M., Philbrick, W., Shulman, G.I., 2008. Muscle-Specific IRS-1 Ser→Ala Transgenic Mice Are Protected From Fat-Induced Insulin Resistance in Skeletal Muscle. Diabetes 57, 2644–2651. https://doi.org/10.2337/db06-0454

O’keefe, M.P., Perez, F.R., Kinnick, T.R., Tischler, M.E., Henriksen, E.J., 2004. Development of whole-body and skeletal muscle insulin resistance after one day of hindlimb suspension. Metabolism. 53, 1215–1222.

Olsen, R.H., Krogh-Madsen, R., Thomsen, C., Booth, F.W., Pedersen, B.K., 2008. Metabolic responses to reduced daily steps in healthy nonexercising men. JAMA 299, 1261–1263. https://doi.org/10.1001/jama.299.11.1259

Pedersen, B.K., 2009. The diseasome of physical inactivity--and the role of myokines in muscle--fat cross talk. J. Physiol. 587, 5559–5568. https://doi.org/10.1113/jphysiol.2009.179515

Perreault, L., Newsom, S.A., Strauss, A., Kerege, A., Kahn, D.E., Harrison, K.A., Snell-Bergeon, J.K., Nemkov, T., D’Alessandro, A., Jackman, M.R., MacLean, P.S., Bergman, B.C., 2018. Intracellular localization of diacylglycerols and sphingolipids influences insulin sensitivity and mitochondrial function in human skeletal muscle. JCI Insight 3. https://doi.org/10.1172/jci.insight.96805

Pierre, N., Appriou, Z., Gratas-Delamarche, A., Derbré, F., 2016. From physical inactivity to immobilization: Dissecting the role of oxidative stress in skeletal muscle insulin resistance and atrophy. Free Radic. Biol. Med. 98, 197–207. https://doi.org/10.1016/j.freeradbiomed.2015.12.028

Rector, R.S., Thyfault, J.P., Laye, M.J., Morris, R.T., Borengasser, S.J., Uptergrove, G.M., Chakravarthy, M.V., Booth, F.W., Ibdah, J.A., 2008. Cessation of daily exercise dramatically alters precursors of hepatic steatosis in Otsuka Long-Evans Tokushima Fatty (OLETF) rats. J. Physiol. 586, 4241–4249. https://doi.org/10.1113/jphysiol.2008.156745

Rector, R.S., Uptergrove, G.M., Borengasser, S.J., Mikus, C.R., Morris, E.M., Naples, S.P., Laye, M.J., Laughlin, M.H., Booth, F.W., Ibdah, J.A., Thyfault, J.P., 2010. Changes in skeletal muscle mitochondria in response to the development of type 2 diabetes or prevention by daily wheel running in hyperphagic OLETF rats. Am. J. Physiol. Endocrinol. Metab. 298, E1179–1187. https://doi.org/10.1152/ajpendo.00703.2009

Reynolds, L.J., Credeur, D.P., Holwerda, S.W., Leidy, H.J., Fadel, P.J., Thyfault, J.P., 2015. Acute inactivity impairs glycemic control but not blood flow to glucose ingestion. Med. Sci. Sports Exerc. 47, 1087–1094. https://doi.org/10.1249/MSS.0000000000000508

Roberts, M.D., Company, J.M., Brown, J.D., Toedebusch, R.G., Padilla, J., Jenkins, N.T., Laughlin, M.H., Booth, F.W., 2012. Potential clinical translation of juvenile rodent inactivity models to study the onset of childhood obesity. Am. J. Physiol. Regul. Integr. Comp. Physiol. 303, R247–258. https://doi.org/10.1152/ajpregu.00167.2012

Salaun, E., Lefeuvre-Orfila, L., Cavey, T., Martin, B., Turlin, B., Ropert, M., Loreal, O., Derbré, F., 2016. Myriocin prevents muscle ceramide accumulation but not muscle fiber atrophy during short-term mechanical unloading. J. Appl. Physiol. Bethesda Md 1985 120, 178–187. https://doi.org/10.1152/japplphysiol.00720.2015

Samuel, V.T., Shulman, G.I., 2012. Mechanisms for insulin resistance: common threads and missing links. Cell 148, 852–871. https://doi.org/10.1016/j.cell.2012.02.017

Sarbassov, D.D., Guertin, D.A., Ali, S.M., Sabatini, D.M., 2005. Phosphorylation and regulation of Akt/PKB by the rictor-mTOR complex. Science 307, 1098–1101. https://doi.org/10.1126/science.1106148

Sargeant, R.J., Pâquet, M.R., 1993. Effect of insulin on the rates of synthesis and degradation of GLUT1 and GLUT4 glucose transporters in 3T3-L1 adipocytes. Biochem. J. 290 (Pt 3), 913–919.

Schmitz-Peiffer, C., 2010. Targeting Ceramide Synthesis to Reverse Insulin Resistance. Diabetes 59, 2351–2353. https://doi.org/10.2337/db10-0912

Singh, B., Saxena, A., 2010. Surrogate markers of insulin resistance: A review. World J. Diabetes 1, 36–47. https://doi.org/10.4239/wjd.v1.i2.36

Softic, S., Kirby, M., Berger, N.G., Shroyer, N.F., Woods, S.C., Kohli, R., 2012. Insulin concentration modulates hepatic lipid accumulation in mice in part via transcriptional regulation of fatty acid transport proteins. PloS One 7, e38952. https://doi.org/10.1371/journal.pone.0038952

Stumvoll, M., Jacob, S., Wahl, H.G., Hauer, B., Löblein, K., Grauer, P., Becker, R., Nielsen, M., Renn, W., Häring, H., 2000. Suppression of systemic, intramuscular, and subcutaneous adipose tissue lipolysis by insulin in humans. J. Clin. Endocrinol. Metab. 85, 3740–3745. https://doi.org/10.1210/jcem.85.10.6898

Tardif, N., Salles, J., Landrier, J.-F., Mothe-Satney, I., Guillet, C., Boue-Vaysse, C., Combaret, L., Giraudet, C., Patrac, V., Bertrand-Michel, J., Migné, C., Chardigny, J.-M., Boirie, Y., Walrand, S., 2011. Oleate-enriched diet improves insulin sensitivity and restores muscle protein synthesis in old rats. Clin. Nutr. Edinb. Scotl. 30, 799–806. https://doi.org/10.1016/j.clnu.2011.05.009

Teich, T., Pivovarov, J.A., Porras, D.P., Dunford, E.C., Riddell, M.C., 2017. Curcumin limits weight gain, adipose tissue growth, and glucose intolerance following the cessation of exercise and caloric restriction in rats. J. Appl. Physiol. Bethesda Md 1985 123, 1625–1634. https://doi.org/10.1152/japplphysiol.01115.2016

Tsujimoto, T., Kajio, H., Sugiyama, T., 2017. Association between hyperinsulinemia and increased risk of cancer death in nonobese and obese people: A population-based observational study. Int. J. Cancer 141, 102–111. https://doi.org/10.1002/ijc.30729

Ussher, J.R., Koves, T.R., Cadete, V.J.J., Zhang, L., Jaswal, J.S., Swyrd, S.J., Lopaschuk, D.G., Proctor, S.D., Keung, W., Muoio, D.M., Lopaschuk, G.D., 2010. Inhibition of De Novo Ceramide Synthesis Reverses Diet-Induced Insulin Resistance and Enhances Whole-Body Oxygen Consumption. Diabetes 59, 2453–2464. https://doi.org/10.2337/db09-1293

Vandesompele, J., De Preter, K., Pattyn, F., Poppe, B., Van Roy, N., De Paepe, A., Speleman, F., 2002. Accurate normalization of real-time quantitative RT-PCR data by geometric averaging of multiple internal control genes. Genome Biol. 3, RESEARCH0034.

Yang, G., Badeanlou, L., Bielawski, J., Roberts, A.J., Hannun, Y.A., Samad, F., 2009. Central role of ceramide biosynthesis in body weight regulation, energy metabolism, and the metabolic syndrome. Am. J. Physiol. Endocrinol. Metab. 297, E211–224. https://doi.org/10.1152/ajpendo.91014.2008

